# Muscle structure governs joint function: linking natural variation in medial gastrocnemius structure with isokinetic plantar flexor function

**DOI:** 10.1101/547042

**Authors:** John F. Drazan, Todd J. Hullfish, Josh R. Baxter

## Abstract

Generating ankle torque is critical for locomotion in elite athletes, the elderly, and many patient populations. Despite the robust findings linking plantar flexor muscle structure to gross function within these populations, the link between variation in plantar flexor fascicle length and ankle kinetics in healthy adults has not been established in the literature. In this study, we determined the relationship between medial gastrocnemius structure and peak torque and total work produced by the plantar flexors during maximal effort contractions. We measured resting fascicle length and pennation angle of the medial gastrocnemius using ultrasound in healthy adult subjects (N=12). Subjects performed maximal effort isometric and isokinetic contractions on a dynamometer. We found that longer fascicles were positively correlated with higher peak torque and total work (R^2^ > 0.41, *p* < 0.013) across all isokinetic velocities, ranging from slow (30 degrees per second) to fast (210 degrees per second) contractions. Higher pennation angles were negatively correlated with peak torque and total work (R^2^ > 0.296, *p* < 0.067). None of these correlations were significant in isometric conditions. To further investigate the coupled effect of fascicle length and pennation angle variation, we used a simple computational model to simulate isokinetic contractions. These simulations confirmed that longer fascicle lengths generate more joint torque and work throughout a greater range of motion. This study provides experimental and computational evidence that links plantar flexor muscle structure and ankle kinetics in healthy young adults, which lends new insight into locomotor function in a range of subpopulations. adults which provides insight into patient recovery following Achilles tendon rupture.

## INTRODUCTION

Plantar flexor function is a critical parameter for human movement in athletes, aging, and patient populations. Comprised of the soleus, lateral gastrocnemius, and medial gastrocnemius; the triceps surae muscles support and accelerate the body during ambulation. Although these plantar flexors appear small relative to knee and hip musculature, plantar flexor function is critical during walking (Graf et al., 2005; Judge et al., 1996), stair climbing (Suzuki et al., 2001), running (Ellis et al., 2014; Nesser et al., 1996), and jumping (Bobbert et al., 1986). Conversely, limited plantar flexor function is associated with decreased walking speed and mobility among elderly populations (Stenroth and Sipila, 2016; Stenroth et al., 2015) and functional deficits in healthy young adults who suffer Achilles tendon injuries (Brorsson et al., 2018). Muscle wasting caused by aging affects the plantar flexors but can be mitigated with resistance training (Morse et al., 2007), highlighting the importance of maintaining muscle structure throughout the lifespan. Of these muscles, the gastrocnemius muscles are particularly important in generating plantar flexor power, due in part to the longer and less pennate muscle fascicles (Lieber and Fridén, 2000).

Gastrocnemius fascicle structure has been linked with plantar flexor function in athletic and patient populations. Trained sprinters have longer gastrocnemius fascicles than non-sprinters and untrained adults (Abe et al., 2000), leading to decreased muscle shortening velocities during a simulated push off of a sprint start (Lee and Piazza, 2009). Even among sprinters, longer and less pennate gastrocnemius fascicles are linked with faster sprint times (Abe et al., 2001; Kumagai et al., 2000). These links between gastrocnemius structure and plantar flexor function translate to patient populations as well. For example, the magnitude of plantar flexor power deficits in patients recovering from Achilles tendon ruptures is strongly correlated with the magnitude of remodeling of the medial gastrocnemius muscle, characterized by shorter resting fascicles (Hullfish et al., 2019a). Longer and less pennate gastrocnemius muscles increase the functional range of ankle motion during simulated muscle contractions by reducing muscle shortening velocity and operating for a longer amount of time in the optimal range of fascicle length (Baxter et al., 2018).

However, the effects of natural variation in gastrocnemius structure on plantar flexor function in healthy adults remains poorly understood. Longer and less pennate muscles operate at slower shortening velocities for a given muscular contraction, explaining the increase amount of muscle force (Lieber and Fridén, 2000). While longer medial gastrocnemius fascicles are correlated with increased muscle shortening speed (Hauraix et al., 2015; Thom et al., 2007), these findings have not been translated to voluntary plantar flexor kinetics measured *in vivo* using isokinetic dynamometry. Given that variation in gastrocnemius muscle structure is well documented (Kawakami et al., 2000) and is modified by injury, (Hullfish et al., 2019a), training (Salzano et al., 2018), and aging (Morse et al., 2005); determining if variations in fascicle length and pennation angle impacts voluntary function has important implications.

The purpose of this study was to determine the relationship between medial gastrocnemius muscle structure and plantar flexor function measured on an isokinetic dynamometer in healthy young adults. Given that longer and less pennate fascicles increase the potential for total muscle shortening velocity, we hypothesized longer fascicles correlate with increased plantar flexor torque and work and that increased pennation would correlate with decreased plantar flexor torque and work during voluntary isokinetic contractions. To test this hypothesis, we quantified medial gastrocnemius fascicle length and pennation angle using ultrasound imaging and measured plantar flexion torque and work in maximal isometric and isokinetic conditions at three rates of ankle rotation on an isokinetic dynamometer. Based on our previous computational modeling (Baxter et al., 2019), we hypothesized that fascicle length would be a stronger correlate of plantar flexor torque and work than pennation angle and muscle thickness. After we correlated medial gastrocnemius structure with plantar flexor isokinetic function, we used a musculoskeletal model to confirm the effects of varying optimal fascicle length and pennation angle on muscle kinetics and shortening velocity.

## METHODS

We quantified medial gastrocnemius structure and plantar flexor function in 12 healthy young adults (6 male, 6 female, age: 25 ± 4.54, BMI: 23.1 ± 4.48) who provided written informed consent in this study which was approved by the University of Pennsylvania IRB (#828374). Subjects were recreationally active and had no reported history of Achilles tendon injury or recent muscle injury in either leg. We acquired measurements of medial gastrocnemius structure and function of the right leg with subjects positioned prone on an isokinetic dynamometer (System 4, Biodex, Shirley, NY). Each subject was positioned on a treatment table that was rigidly secured to the dynamometer while wearing standardized lab shoes. After their foot was secured to the foot plate with the medial malleolus aligned with the spindle of the dynamometer, each subject selected their ankle range of motion by fully dorsiflexing and plantar flexing their ankle. Once the subject specific range of motion was set, the investigator set the ankle neutral position, which was recorded for post-processing. To ensure consistency between subjects, all experimental procedures were performed by a single investigator.

To quantify medial gastrocnemius structure, we acquired ultrasound images of the medial gastrocnemius throughout the entire passive range of motion of each subject. We positioned the 7.5 MHz, 6 cm probe (LV7.5/60/128Z-2, SmartUs, TELEMED) over the mid-substance of the muscle belly and secured it to the leg using a custom-made cast and strap system. Ultrasound images were acquired between 30 and 60 Hz while each subject’s ankle was moved through its passive range of motion moving from plantar flexion to dorsiflexion at a rate of 10 degrees/s. We quantified resting fascicle length, pennation angle, and muscle thickness with the ankle at the resting angle rather than at neutral to avoid stretching the fascicles (Aeles et al., 2017). This resting angle was set to 16 degrees plantar flexion for all subjects based on values of average resting ankle position across 42 patients reported in the literature (Zellers et al., 2018). We manually identified the superficial and deep aponeuroses and a muscle fascicle by selecting two points on each structure to form a line (**Figure 1**). This manual approach of fascicle measurement has been demonstrated to be repeatable within the same observer (Drazan et al., 2019). We calculated fascicle length as the distance between the superficial and deep attachments of the fascicle and pennation angle as the angle between the fascicle and the deep aponeurosis. Muscle thickness was calculated as the fascicle length multiplied by the sine of the pennation angle. We measured fascicles that were in the middle of the imaging frame for all subjects. In case that the entire fascicle was not in frame, linear extrapolation was used to calculate fascicle length. We averaged each measurement of structure across three passive range of motion trials to determine the measures of resting muscle structure we used in our analyses.

**Figure 1 (1 column).**
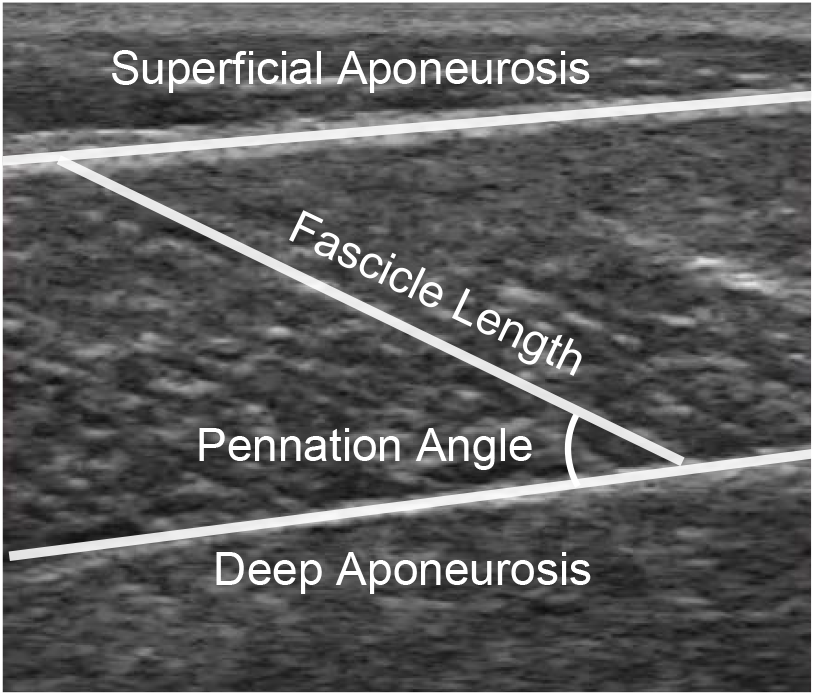
Resting muscle architecture was measured from an ultrasound frame captured at 16° plantar flexion. An examiner identified the two aponeuroses as well as a fascicle within the frame, each of which was approximated as a straight line. Pennation angle was calculated as the angle between the fascicle and the deep aponeurosis and muscle thickness was calculated as the sine of the pennation angle times the length of the fascicle.

To determine plantar flexor work and peak torque, subjects performed maximal voluntary plantar flexor contractions on an isokinetic dynamometer. Subjects performed maximal isometric contractions at neutral ankle angle and isokinetic contractions at three rotational velocities across their range of motion. The subject’s right foot was secured to a foot plate with the medial malleolus of the ankle aligned with the dynamometer’s spindle. First, we measured peak isometric plantar flexor torque with the ankle at neutral with subjects in prone position. Next, peak isokinetic plantar flexion contractions were performed throughout a subject’s entire range of motion at three speeds: slow (30 degrees per second), medium (120 degrees per second), and fast (210 degrees per second). At the start of each of these trials, we confirmed that the subject positioned their ankle in their peak dorsi-flexion angle. We provided verbal encouragement (McNair et al., 1996) as well as visual feedback to ensure that subjects maximally contract their plantar flexors during each condition. Contractions were not ramped, instead subjects were instructed to immediately “push as hard and as fast as they could.” Subjects continued to perform maximal plantar flexion contractions for each test condition until the peak torque was consistent for two consecutive trials.

To test our hypothesis that fascicle length would be positively and strongly correlated with plantar flexor function and that pennation angle would be negatively and moderately correlated with plantar flexor function, we performed univariate linear regression to determine the relationship between measures of medial gastrocnemius structure with peak plantar flexor torque and work at each contraction condition. We calculated the strength correlation for each of these regression analyses using the coefficient of determination (R^2^), which is an index of the correlation strength ranging between 0 and 1 where values between 0 and 0.04 indicate negligible correlation, 0.04 and 0.25 indicate weak correlation, 0.25 and 0.64 indicate moderate correlation, and 0.64 and 1 indicate strong correlation (Morton et al., 2005). We set an *a priori* alpha level of 0.05 and performed all statistical analysis using scientific computing software (MATLAB, MathWorks, Natick, MA).

Because muscle structure and anthropometry varied in our subjects, we quantified muscle thickness and lower-leg length to determine if these factors accounted for some of the variability in plantar flexor kinetics. We calculated muscle thickness as the product of the resting fascicle length and the sine of the pennation angle. We quantified lower-leg length by measuring the distance between reflective markers placed on the lateral malleolus and the proximal head of the fibula that was measured using a 12 camera motion capture system (Raptor Series, Motion Analysis Corporation, Rohnert Park, CA). To evaluate the effect of patient stature on resting muscle structure, we linearly regressed fascicle length, pennation angle, and muscle thickness against body mass and leg length. Additionally, muscle thickness was linearly regressed against fascicle length and pennation angle.

We simulated the 3 isokinetic contraction speeds to demonstrate how variations in fascicle length and pennation angle affected joint kinetics and muscle shortening velocity (Fig. 4A) using a musculoskeletal model (Baxter and Hast, 2019; Delp et al., 2007). Briefly, the ankle was constrained by a pin joint and actuated by a combined gastrocnemius muscle, soleus muscle, and tibialis anterior muscle. The musculoskeletal model was positioned in the prone position and resting ankle angle was set at 16 degrees plantar flexion, which matched previous literature reports (Zellers et al., 2018) and the ankle angle at which we measured medial gastrocnemius structure in the current study. During each test speed, we changed the optimal fascicle length (which we will refer to as fascicle length for consistency with our *in vivo* measurements) and pennation angle of both the gastrocnemius and soleus muscles. We set the fascicle length of the gastrocnemius muscle to 64 mm and the pennation angle to 22 degrees, which we acquired from ultrasound images of our test subjects in 16 degrees of plantar flexion. The soleus muscle was set to the model default values of 44 mm for fascicle length and 28 degrees of pennation. We iteratively adjusted by fascicle length and pennation angle by 10%, ranging from 50% to 150% of the model default values (Fig. 4A). During each test iteration, we used a gradient based optimization procedure to find the tendon slack lengths that placed the ankle in static equilibrium. We then simulated maximal plantar flexor contractions at 30, 120, and 210 degrees/s and recorded the muscle force generated and shortening velocities of the gastrocnemius and soleus muscles. To test the effects of variations in gastrocnemius structure, we analyzed the contributions of gastrocnemius force towards ankle kinetics (complete model and simulation results available in supplemental data).

## RESULTS

Resting fascicle length was positively and moderately correlated with plantar flexor work (**Figure 2A**) and peak torque (**Figure 3A**) during isokinetic plantar flexion contractions. More than half of the variability in plantar flexor work (R^2^ = 0.599, *P* = 0.003) and peak torque (R^2^ = 0.521, *P* = 0.008) during 30°/s isokinetic contractions was explained by resting fascicle length. However, the correlation between resting fascicle length and plantar flexor work and peak torque decreased during faster isokinetic contractions at 120°/s (0.413 > R^2^ > 0.415, *p* < 0.024) and 210°/s (0.477 > R^2^ > 0.494, *P* < 0.013). Fascicle length had the weakest correlation with peak torque during isometric conditions (R^2^ = 0.325, *P* = 0.053). Subjects generated less torque and did less work as rotational velocity increased (**Table 1**).

**Figure 2 (1 column).**
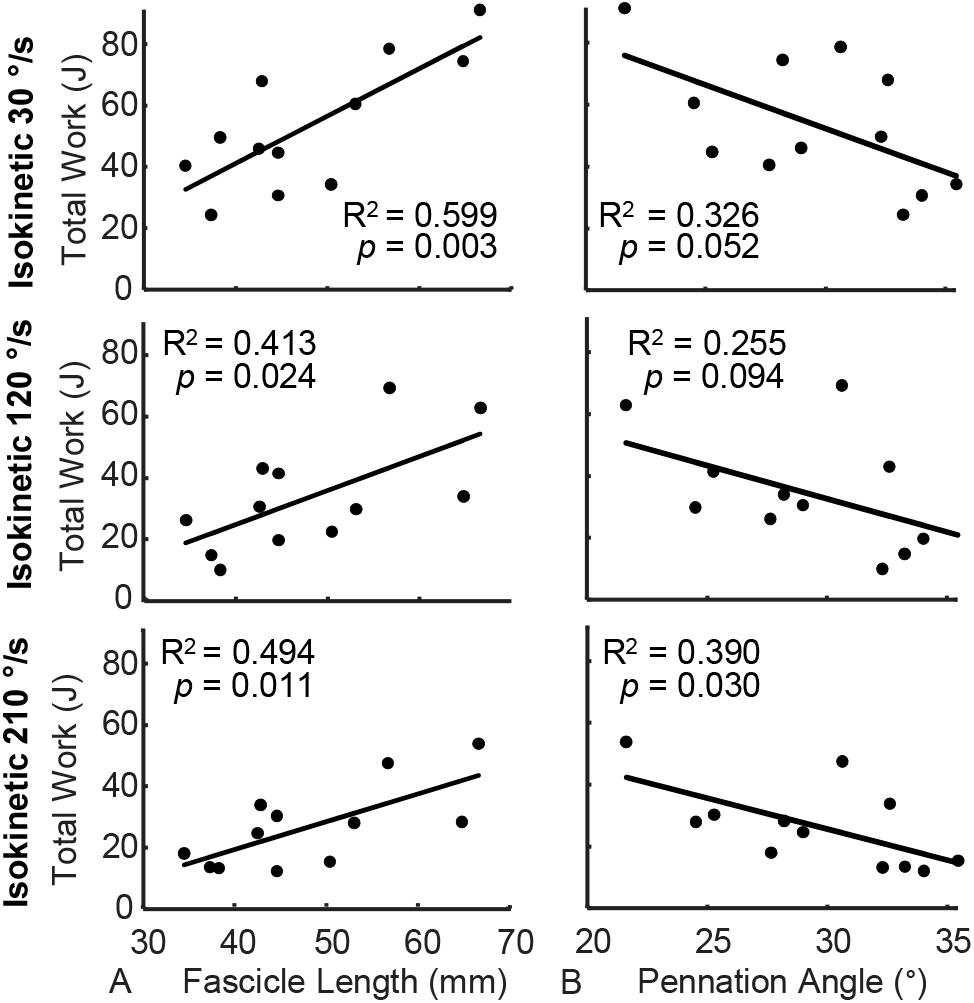
Plantar flexor work positively correlated with resting fascicle length during maximal isokinetic contractions. Plantar flexor work produced at three isokinetic speeds (30°/s – top row, 120°/s – middle row, and 210°/s – bottom row) positively correlated with (A) fascicle length and negatively correlated with (B) pennation angle. Fascicle lengths explained more variation in plantar flexor work than pennation angle (N=12).

**Figure 3 (1 column).**
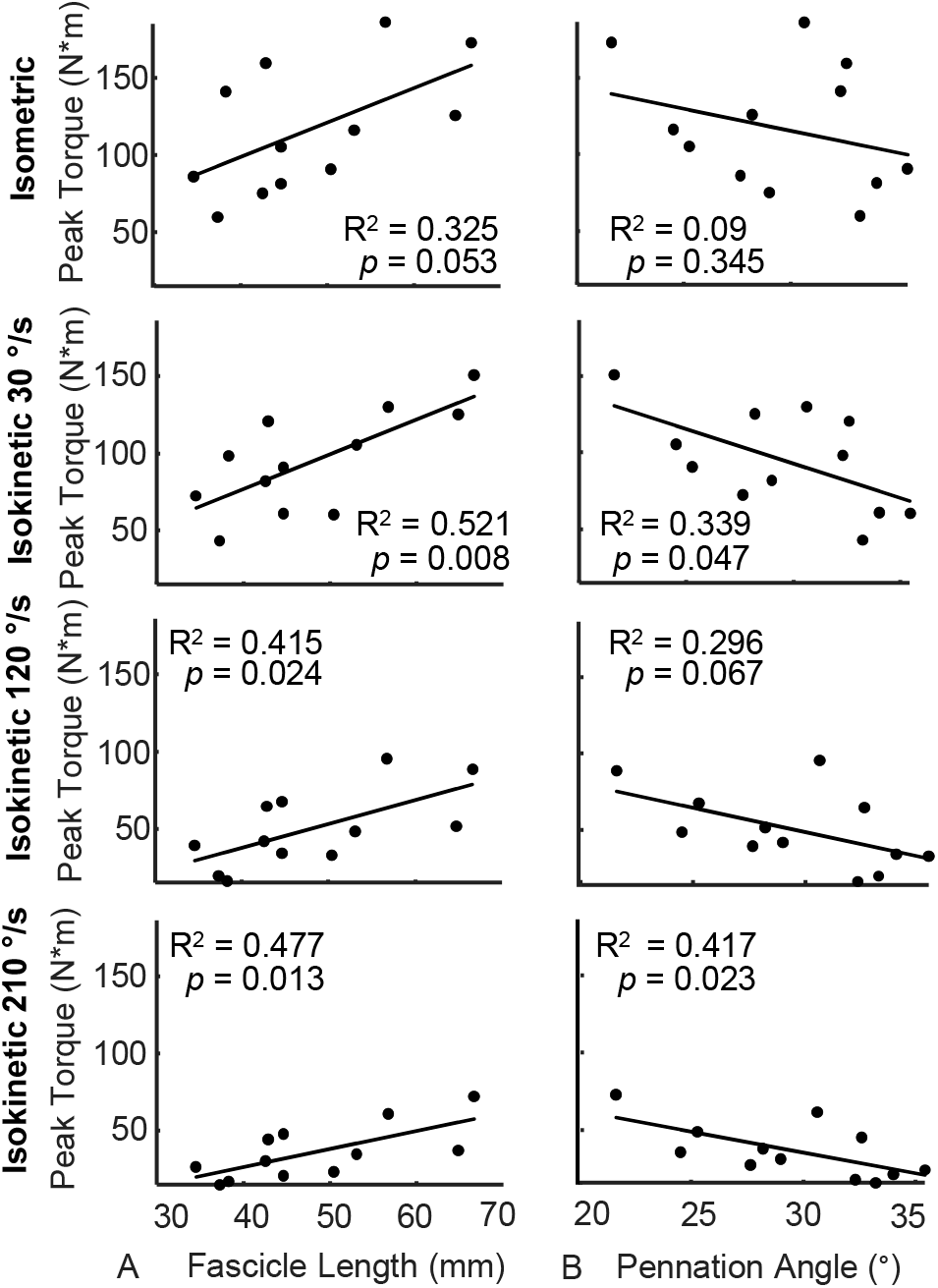
Peak plantar flexor torque positively correlated with resting fascicle length during maximal isokinetic contractions. Peak plantar flexor torque produced at three isokinetic speeds (30°/s – second row, 120°/s – third row, and 210°/s – bottom row) is positively correlated with (A) fascicle length and negatively correlated with (B) pennation angle. Conversely, fascicle length explains less variation in peak isometric torque (top row) and pennation angle is not correlated with isometric torque. Fascicle lengths explained more variation in plantar flexor work than pennation angle (N=12).

**Table 1.** Descriptive data on subject plantar flexor function and musculoskeletal parameters.

| | Mean ± Standard Deviation |
| --- | --- |
| **Plantarflexor Kinetics** |  |
| Peak Isometric Torque (N*m) | 116.8 ± 39.1 |
| Peak Isokinetic Torque $30^\circ/\text{s}$ (N*m) | 95.1 ± 31.3 |
| Peak Isokinetic Torque $120^\circ/\text{s}$ (N*m) | 50.1 ± 24.0 |
| Peak Isokinetic Torque $210^\circ/\text{s}$ (N*m) | 35.9 ± 17.0 |
| Total Work $30^\circ/\text{s}$ (J) | 53.7 ± 20.0 |
| Total Work $120^\circ/\text{s}$ (J) | 33.8 ± 17.3 |
| Total Work $210^\circ/\text{s}$ (J) | 26.6 ± 12.9 |
| **Musculoskeletal Parameters** |  |
| Range of Motion (degrees) | 53.7 ± 4.2 |
| Fascicle Length (mm) | 48.0 ± 10.0 |
| Pennation Angle (degrees) | 29.5 ± 4.1 |
| Muscle Thickness (mm) | 23.2 ± 4.2 |
| Leg Length (mm) | 362.2 ± 36.9 |
| Mass (kg) | 67.3 ± 19.3 |

Pennation angle was negatively and moderately correlated with plantar flexor work (0.255 > R^2^ > 0.39, *p* < 0.052) and peak torque (0.296 > R^2^ > 0.417, *p* < 0.067 during isokinetic plantar flexion contractions (**Figure 2B and 3B**). However, these correlations were weaker than resting fascicle length for each test condition and only reached statistical significance during 30°/s (*P* = 0.047) and 210 °/s (*P* = 0.023) conditions for measurements of peak torque. Peak isometric torque was not explained by resting pennation angle (R^2^ = 0.09, *P* = 0.345).

Muscle thickness was moderately and significantly correlated with fascicle length (R^2^ = 0.536, *p* < 0.001) but not correlated with pennation angle (R^2^ = 0.064, *P* = 0.084). Despite the positive correlation between resting muscle thickness and fascicle length, resting muscle thickness was not significantly correlated with function at any isokinetic testing condition (**Table 2**). Muscle thickness was positively and weakly correlated with peak torque at isometric max (R^2^ = 0.161). However, this correlation did not reach statistical significance (*P* = 0.197).

**Table 2.** Correlations between muscle thickness and plantarflexion kinetics

|  | Peak Torque | Total Work |
| --- | --- | --- |
| Isometric Max | 0.153<br>(0.209) | - |
| Isokinetic 30 °/s | 0.124<br>(0.262) | 0.172<br>(0.181) |
| Isokinetic 120 °/s | 0.104<br>(0.307) | 0.117<br>(0.276) |
| Isokinetic 210 °/s | 0.076<br>(0.387) | 0.089<br>(0.347) |

Correlations of determination ( $R^2$ ) and regression $p$ -values are reported as $R^2$ ( $p$ -value)

Muscle structure was weakly correlated with subject stature (**Table 3**). Fascicle length was weakly correlated with leg length (R^2^ = 0.084, P = 0.046) and body mass (R^2^ = 0.132, *P* = 0.011). Pennation angle was weakly correlated with body mass (R^2^ = 0.098, *P* = 0.030) but not leg length. Muscle thickness was not correlated with either leg length or body mass.

**Table 3.** Correlations between muscle structure and subject stature.

|  | Leg Length | Mass | Muscle Thickness |
| --- | --- | --- | --- |
| Fascicle Length | 0.084<br>(0.046) | 0.132<br>(0.011) | 0.53<br>($<0.001$) |
| Pennation Angle | 0.013<br>(0.445) | 0.098<br>(0.030) | 0.063<br>(0.083) |
| Muscle Thickness | 0.070<br>(0.069) | 0.037<br>(0.193) | - |

Correlations of determination ( $R^2$ ) and regression $p$ -values are reported as $R^2$ ( $p$ -value)

Longer muscle fascicles had a greater effect on simulated plantar flexor function compared to similar decreases in pennation angle (**Figure 4B and C**). The effects of small increases in fascicle length increased with greater rates of ankle rotation during these simulated isokinetic plantar flexion contractions. A 1 percent increase in the gastrocnemius fascicle length led to a 0.3% increase in peak plantar flexor torque at 30 degrees per second and 0.8% increase in peak plantar flexor torque at 210 degrees per second (**Figure 4B**). These small increases in gastrocnemius fascicle length had a greater effect on simulated plantar flexor work done by the ankle joint (**Figure 4C**). Increasing the gastrocnemius fascicle length by 1% increased joint work by 0.6% during 30 degrees per second contractions and 1.0% during 210 degrees per second contractions.

**Figure 4 (2 column).**
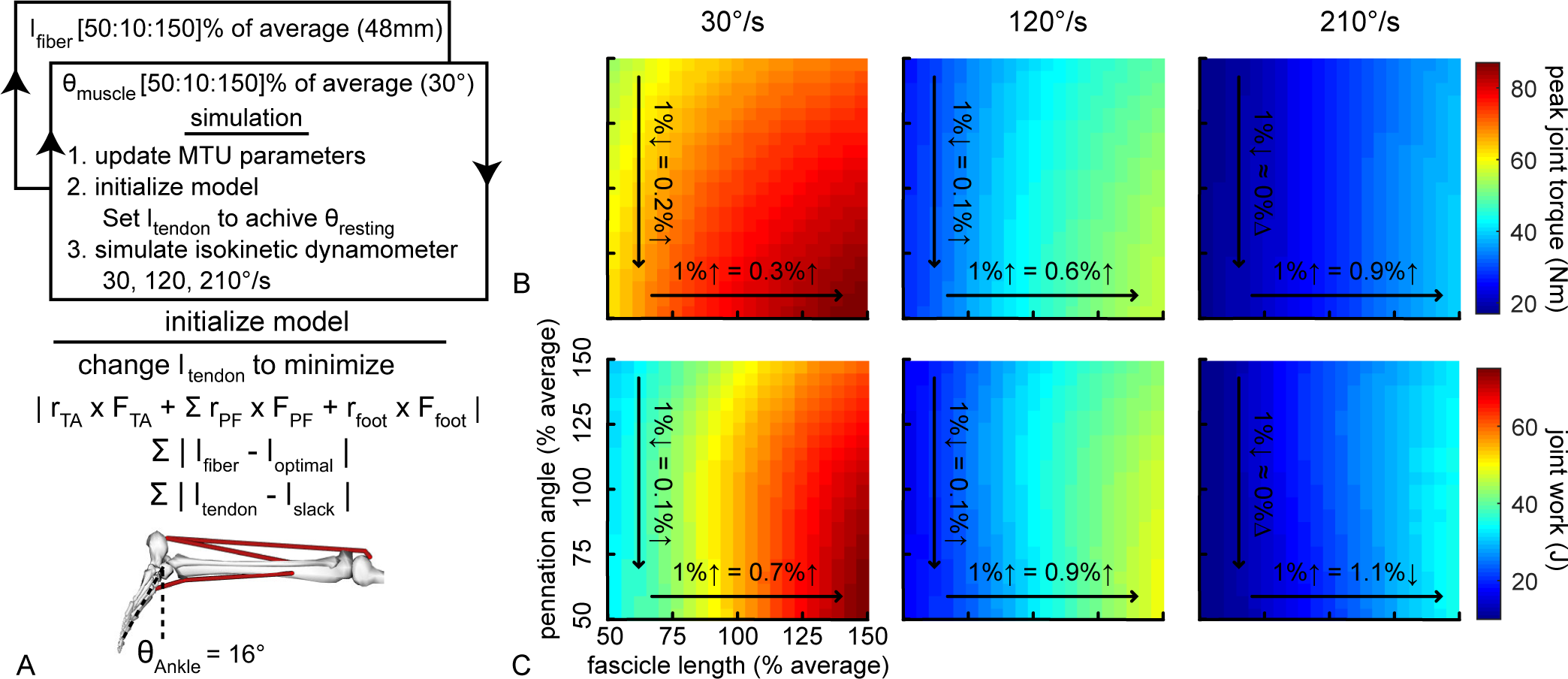
Longer muscle fascicles generate greater plantar flexor torque and work during simulated isokineticcontractions. Wesimulated the effects of varying fascicle length (*x-axis*) and pennation angle (*y-axis*) using a simplified musculoskeletal model of the lower leg (A). After each model permutation was initialized by solving for tendon slack lengths to reach static equilibrium at 16 degrees plantar flexion, we simulated maximal isokinetic plantar flexion contractions at 30, 120, and 210 degrees per second. Small increases in fascicle length had a greater effect on peak torque (B) than similarly small decreases in pennation angle. Similar to peak joint torque, longer fascicles also increased the amount of work done by the muscle during maximal contractions at each on speed (C). We tested a wide range (± 50%) of fascicle lengths and pennation angles centered at the average measurements made in the *in vivo* experiment. Peak torque was calculated as the product of the active contributions of the gastrocnemius muscle and the muscle moment arm (Nm). Joint work was calculated as the integral of the torque-angle curve.

These increases in plantar flexor kinetics caused by longer gastrocnemius muscle fascicles were explained by 2 factors (**Figure 5**). First, the longer muscle fascicles generated greater force at each joint angle. Second, the longer muscle fascicles continued to generate muscle force in deeper plantar flexion while the shorter muscle fascicles stopped producing force starting at 40 degrees of plantar flexion. However, similar changes in pennation angle had weaker effects on plantar flexor kinetics (**Figure 4 B and C**).

**Figure 5 (1 column).**
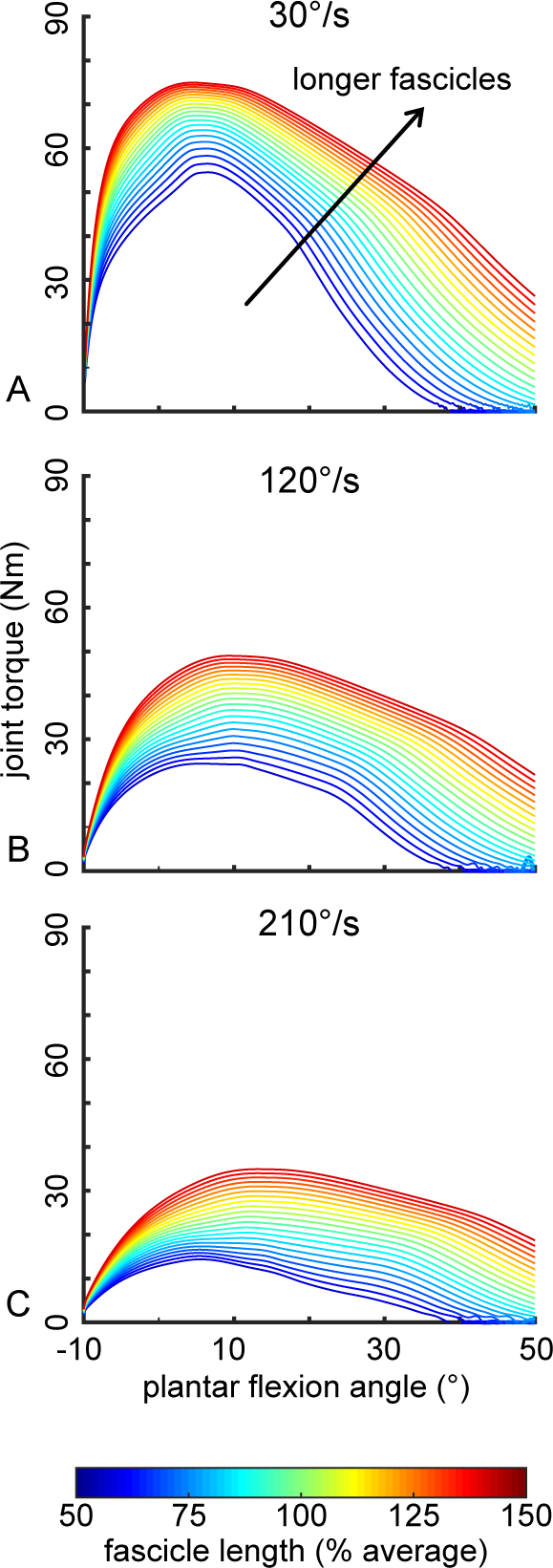
Longer muscle fascicles lead to increased plantar flexor torque throughout the ankle range of motion during simulated isokinetic contractions. Longer gastrocnemius fascicles (more red) generated greater amounts of plantar flexor torque throughout the entire range of motion compared to shorter fascicles (more blue). These effects were similar at slow (A), medium (B), and fast (C) simulated isokinetic contraction speeds.

## DISCUSSION

In this study, we demonstrated the relationship between plantar flexor structure and function in a cohort of healthy young adults. Our findings support our hypothesis that resting fascicle length is positively correlated with peak plantar flexor torque and work while pennation angle has a smaller, negative correlation with peak plantar flexor torque and work. To determine if plantarflexor function was simply explained by muscle thickness, we regressed muscle thickness with plantar flexor kinetics and found no effect on plantar flexor torque or work in any condition. Although fascicle length has been positively correlated with muscle force in isolated muscle experiments (Lieber, 1997); to our knowledge, this is the first study to link natural variation in gastrocnemius fascicle length with plantar flexor torque and work in healthy adults (Ema et al., 2016).

Our measurements of medial gastrocnemius structure and isokinetic plantar flexor torque and work compare favorably with previous reports in the literature. We decided to measure resting muscle structure at 16 degrees plantar flexion to approximate the ankle angle at which medial gastrocnemius muscle-tendon slack occurs (Zellers et al., 2018). Other measurements of gastrocnemius structure were acquired with the ankle either under load or neutrally aligned which may explain slightly longer (4-9 mm) and less pennate (6-11 degrees) muscle fascicles (Baxter and Piazza, 2014; Kubo et al., 2003; Thom et al., 2007) than those we measured in this current study. Similarly, our measurements of pennation at slack are comparable to earlier measurements in the medial gastrocnemius at the same position (Hoang et al., 2007). Our measurements of plantar flexion torque and work capacity compares well with the literature. Our values for torque and work done at 30 and 120 degrees per second compares well with previously reported values (Randhawa and Wakeling, 2013; Woodson et al., 1995) and our values for maximal isometric torque compares well with studies with similar subject positioning (Arampatzis et al., 2006). A previous study reported higher values for torque generation, however in this case, the subjects were all male and were seated rather than positioned in prone (Baxter and Piazza, 2014).

Our results suggest that fascicle length and pennation angle, which govern absolute muscle shortening velocity (Lieber and Ward, 2011), have greater effects on isokinetic plantar flexor function than muscle thickness. Our results are consistent with the force-length and force-velocity properties of muscle. While fascicles of different lengths undergo the same absolute contractile velocity during a given isokinetic contraction, longer fascicles have additional sarcomeres in series which extend the functional operating length of the muscle while reducing relative shortening velocity. This enables longer fascicles to operate at slower velocities on the force-velocity curve, increasing force production at all isokinetic speeds (**Figure 6A**) (Lieber and Fridén, 2000). Similarly, the force-velocity effects are also affected by variation in pennation angle. Greater muscle pennation increases the fascicle shortening demands for a given muscle shortening contraction. Thus, more pennate muscles generate less force and do less work in isokinetic conditions (**Figure 6B**). Our experimental findings that fascicle length explained more variation in plantar flexor kinetics than pennation angle agrees with the theoretical framework of the force-velocity properties of muscle and our recent computational model (Baxter et al., 2019) as well as the computational results described in this paper (**Figure 4 and 5**).

**Figure 6 (1 column).**
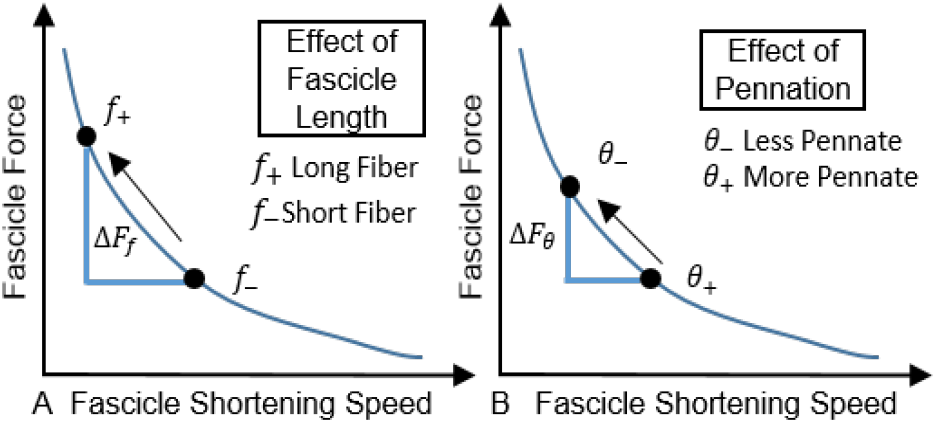
Longer and less pennate muscle fascicles reduce the shortening speed demands during a given isokinetic contraction. Longer and (A) less pennate (B) fascicles decrease relative shortening velocity during a given joint rotation, which results in increased fascicle force. The force-velocity properties of skeletal muscle are more sensitive to changes in fascicle length than pennation angle, which agrees with our experimental findings that ankle torque is more strongly correlated with fascicle length than pennation angle.

Our results highlight the effects of natural variability in muscle fascicle lengths on plantar flexor function. Force-length characteristics have been demonstrated in humans *in vivo* (Maganaris, 2001), however these results were only reported in isometric contractions. Despite evidence of variations in muscle structure between different populations of human subjects (Abe et al., 2000; Hullfish et al., 2019b; Kearns et al., 2000; Kubo et al., 2003; Lee and Piazza, 2009), there are few studies reporting the relationship between variations in resting fascicle length or pennation angle and dynamic muscle function in humans (Ema et al., 2016). Outside of the plantar flexors, one study found that elbow extensor velocity under no load was positively correlated with muscle volume and pennation angle but not muscle length (Wakahara et al., 2013). However, this previous study did not directly measurement fascicle length, instead they used muscle length as a surrogate for fascicle length. Differences in fascicle length explains almost 50% of reduced fascicle shortening velocity between young and old men (Thom et al., 2007). Our previous work did not find a relationship between fascicle length and peak torque during isokinetic contractions at 30, 120 and 210 °/s. However, these previous studies measured fascicle length at neutral, which is a less reliable position to measure fascicle length given the sensitive relationship between passive fascicle load and length (Aeles et al., 2017).

We decided to measure the medial gastrocnemius because of previous observations linking structural changes in that muscle following Achilles tendon rupture to functional deficits (Baxter et al., 2018; Hullfish et al., 2019a) coupled with similar observations with other groups (Peng et al., 2017; Peng et al., 2019). Prior groups linking plantar flexor torque measurements with medial gastrocnemius structure without accounting for the other two triceps surae muscles (Hauraix et al., 2015; Thom et al., 2007). These groups approximated the torque generated by the gastrocnemius by multiplying total plantar flexor torque values by constant value ranging from 0.159 (Fukunaga et al., 1996) to 0.218 (Morse et al., 2005). As these are constant values, this adjustment would not affect the correlations we have found in this study. In addition, we performed an analysis on the publically available data set (Crouzier et al., 2018) and determined that fascicle length and pennation angles of the medial gastrocnemius both correlate with the lateral gastrocnemius and soleus muscles (R^2^ > 0.413, *P* < 0.023).

This study was affected by several limitations. Despite having a relatively small sample size (N=12), our *in vivo* findings agree with both basic principles of skeletal muscle mechanics (Lieber and Fridén, 2000) and our computational model. We did not confirm maximal effort during contractions across the triceps surae using electromyography. Instead, we provided subjects verbal encouragement and real-time visual feedback (Drazan et al., 2019) and had each subject repeat each maximal contraction condition until their peak torques were consistent between consecutive trials (McNair et al., 1996). We used muscle thickness as a proxy for muscle volume, which is positively correlated with measurements of muscle volume acquired with magnetic resonance imaging (R^2^ = 0.527, *P* < 0.001) (Crouzier et al., 2018) and cadaveric measurements (R^2^ = 0.497, *P* = 0.017) (Bandholm et al., 2007). We did not quantify plantar flexor moment arm, which may affect muscle shortening dynamics and plantar flexor kinetics (Baxter and Piazza, 2014; Nagano and Komura, 2003). While considering medial gastrocnemius structure to be a surrogate measure of the triceps surae muscles (Crouzier et al., 2018), variations in lateral gastrocnemius and soleus structure might strengthen the correlation between plantar flexor structure and function, it will not decrease our observed correlations.

In conclusion, our study demonstrates the link between resting structure of the medial gastrocnemius with isokinetic plantar flexion function. These findings may have important implications on plantar flexor function following muscle remodeling elicited through injury, training, and aging. However, the link between isokinetic plantar flexor function and ambulatory function requires further investigation, and future work should directly test the link between muscle structure and movement biomechanics.

## Acknowledgements

We would like to thank R. Mathew and S. Donde for assistance with data collection and processing.

## REFERENCES

Abe, T., Kumagai, K. and Brechue, W. F. (2000). Fascicle length of leg muscles is greater in sprinters than distance runners. Med. Sci. Sports Exerc. 32, 1125–1129.

Abe, T., Fukashiro, S., Harada, Y. and Kawamoto, K. (2001). Relationship between sprint performance and muscle fascicle length in female sprinters. J. Physiol. Anthropol. Appl. Human Sci. 20, 141–147.

Aeles, J., Lenchant, S., Vanlommel, L. and Vanwanseele, B. (2017). Bilateral differences in muscle fascicle architecture are not related to the preferred leg in jumping athletes. Eur. J. Appl. Physiol. 117, 1453–1461.

Arampatzis, A., Karamanidis, K., Stafilidis, S., Morey-Klapsing, G., DeMonte, G. and Brüggemann, G.-P. (2006). Effect of different ankle-and knee-joint positions on gastrocnemius medialis fascicle length and EMG activity during isometric plantar flexion. J. Biomech. 39, 1891–1902.

Bandholm, T., Sonne-Holm, S., Thomsen, C., Bencke, J., Pedersen, S. A. and Jensen, B. R. (2007). Calf Muscle Volume Estimates: Implications for Botulinum Toxin Treatment? Pediatr. Neurol. 37, 263–269.

Baxter, J. R. and Hast, M. W. (2019). Plantarflexor metabolics are sensitive to resting ankle angle and optimal fiber length in computational simulations of gait. Gait Posture 67, 194–200.

Baxter, J. R. and Piazza, S. J. (2014). Plantar flexor moment arm and muscle volume predict torque-generating capacity in young men. J. Appl. Physiol. 116, 538–44.

Baxter, J. R., Hullfish, T. J. and Chao, W. (2018). Functional deficits may be explained by plantarflexor remodeling following Achilles tendon rupture repair: Preliminary findings. J. Biomech. 79, 238–242.

Baxter, J. R., Farber, D. C. and Hast, M. W. (2019). Plantarflexor fiber and tendon slack length are strong determinates of simulated single-leg heel raise height. J. Biomech.

Bobbert, M. F., Huijing, P. A. and van Ingen Schenau, G. J. (1986). An estimation of power output and work done by the human triceps surae musle-tendon complex in jumping. J. Biomech. 19, 899–906.

Brorsson, A., Grävare Silbernagel, K., Olsson, N. and Nilsson Helander, K. (2018). Calf Muscle Performance Deficits Remain 7 Years After an Achilles Tendon Rupture. Am. J. Sports Med. 46, 470–477.

Crouzier, M., Lacourpaille, L., Nordez, A., Tucker, K. and Hug, F. (2018). Neuromechanical coupling within the human triceps surae and its consequence on individual force-sharing strategies. J. Exp. Biol. 221, jeb187260.

Delp, S. L., Anderson, F. C., Arnold, A. S., Loan, P., Habib, A., John, C. T., Guendelman, E. and Thelen, D. G. (2007). OpenSim: open-source software to create and analyze dynamic simulations of movement. IEEE Trans. Biomed. Eng. 54, 1940–1950.

Drazan, J. F., Hullfish, T. J. and Baxter, J. R. (2019). An automatic fascicle tracking algorithm quantifying gastrocnemius architecture during maximal effort contractions. PeerJ 7, e7120.

Ellis, R. G., Sumner, B. J. and Kram, R. (2014). Muscle contributions to propulsion and braking during walking and running: Insight from external force perturbations. Gait Posture 40, 594–599.

Ema, R., Akagi, R., Wakahara, T. and Kawakami, Y. (2016). Training-induced changes in architecture of human skeletal muscles: Current evidence and unresolved issues. J. Phys. Fit. Sports Med. 5, 37–46.

Fukunaga, T., Roy, R. R., Shellock, F. G., Hodgson, J. A. and Edgerton, V. R. (1996). Specific tension of human plantar flexors and dorsiflexors. J. Appl. Physiol. Bethesda Md 1985 80, 158–165.

Graf, A., Judge, J. O., Õunpuu, S. and Thelen, D. G. (2005). The effect of walking speed on lower-extremity joint powers among elderly adults who exhibit low physical performance. Arch. Phys. Med. Rehabil. 86, 2177–2183.

Hauraix, H., Nordez, A., Guilhem, G., Rabita, G. and Dorel, S. (2015). In vivo maximal fascicle-shortening velocity during plantar flexion in humans. J. Appl. Physiol. 119, 1262–1271.

Hoang, P. D., Herbert, R. D., Todd, G., Gorman, R. B. and Gandevia, S. C. (2007). Passive mechanical properties of human gastrocnemius muscle tendon units, muscle fascicles and tendons in vivo. J. Exp. Biol. 210, 4159–4168.

Hullfish, T. J., O’Connor, K. M. and Baxter, J. R. (2019a). Gastrocnemius muscle remodeling explains functional deficits three months following Achilles tendon rupture. BioRxiv Prepr. 585505.

Hullfish, T. J., O’Connor, K. M. and Baxter, J. R. (2019b). Gastrocnemius fascicles are shorter and more pennate throughout the first month following acute Achilles tendon rupture. PeerJ 7, e6788.

Judge, J. O., Davis, R. B. and Ounpuu, S. (1996). Step length reductions in advanced age: the role of ankle and hip kinetics. J. Gerontol. A. Biol. Sci. Med. Sci. 51, M303–M312.

Kawakami, Y., Ichinose, Y., Kubo, K., Ito, M., Imai, M. and Fukunaga, T. (2000). Architecture of Contracting Human Muscles and Its Functional Significance. J. Appl. Biomech. 16, 88–97.

Kearns, C. F., Abe, T. and Brechue, W. F. (2000). Muscle enlargement in sumo wrestlers includes increased muscle fascicle length. Eur. J. Appl. Physiol. 83, 289–296.

Kubo, K., Kanehisa, H., Azuma, K., Ishizu, M., Kuno, S.-Y., Okada, M. and Fukunaga, T. (2003). Muscle Architectural Characteristics in Young and Elderly Men and Women. Int. J. Sports Med. 24, 125–130.

Kumagai, K., Abe, T., Brechue, W. F. W. F., Ryushi, T., Takano, S. and Mizuno, M. (2000). Sprint performance is related to muscle fascicle length in male 100-m sprinters. J. Appl. Physiol. 88, 811–816.

Lee, S. S. M. and Piazza, S. J. (2009). Built for speed: musculoskeletal structure and sprinting ability. J. Exp. Biol. 212, 3700–3707.

Lieber, R. L. (1997). Muscle fiber length and moment arm coordination during dorsi-and plantarflexion in the mouse hindlimb. Acta Anat. (Basel) 159, 84–89.

Lieber, R. L. and Fridén, J. (2000). Functional and clinical significance of skeletal muscle architecture. Muscle Nerve 23, 1647–1666.

Lieber, R. L. and Ward, S. R. (2011). Skeletal muscle design to meet functional demands. Philos. Trans. R. Soc. Lond. B. Biol. Sci. 366, 1466–1476.

Maganaris, C. N. C. C. N. (2001). Force-length characteristics of in vivo human skeletal muscle. Acta Physiol Scand 172, 279–285.

McNair, P. J., Depledge, J., Brettkelly, M. and Stanley, S. N. (1996). Verbal encouragement: effects on maximum effort voluntary muscle: action. Br. J. Sports Med. 30, 243–245.

Morse, C. I., Thom, J. M., Birch, K. M. and Narici, M. V. (2005). Changes in triceps surae muscle architecture with sarcopenia. Acta Physiol. Scand. 183, 291–298.

Morse, C. I., Thom, J. M., Mian, O. S., Birch, K. M. and Narici, M. V. (2007). Gastrocnemius specific force is increased in elderly males following a 12-month physical training programme. Eur. J. Appl. Physiol. 100, 563–570.

Morton, R. F., Hebel, J. R. and McCarter, R. J. (2005). A Study Guide to Epidemiology and Biostatistics. Jones & Bartlett Learning.

Nagano, A. and Komura, T. (2003). Longer moment arm results in smaller joint moment development, power and work outputs in fast motions. J. Biomech. 36, 1675–1681.

Nesser, T. W., Latin, R. W., Berg, K. and Prentice, E. (1996). Physiological Determinants of 40-Meter Sprint Performance in Young Male Athletes: J. Strength Cond. Res. 10, 263–267.

Peng, W.-C., Chang, Y.-P., Chao, Y.-H., Fu, S., Rolf, C., Shih, T. T., Su, S.-C. and Wang, H.-K. (2017). Morphomechanical alterations in the medial gastrocnemius muscle in patients with a repaired Achilles tendon: Associations with outcome measures. Clin. Biomech. 43, 50–57.

Peng, W.-C., Chao, Y.-H., Fu, A. S. N., Fong, S. S. M., Rolf, C., Chiang, H., Chen, S. and Wang, H.-K. (2019). Muscular Morphomechanical Characteristics After an Achilles Repair. Foot Ankle Int. 1071100718822537.

Randhawa, A. and Wakeling, J. M. (2013). Associations between muscle structure and contractile performance in seniors. Clin. Biomech. 28, 705–711.

Salzano, M. Q., Cox, S. M., Piazza, S. J. and Rubenson, J. (2018). American Society of Biomechanics Journal of Biomechanics Award 2017: High-acceleration training during growth increases optimal muscle fascicle lengths in an avian bipedal model. J. Biomech. 80, 1–7.

Stenroth, L. and Sipila, S. (2016). Slower Walking Speed in Older Men Improves Triceps Surae Force Generation Ability. Med. Sci. Sports Exerc.

Stenroth, L., Sillanpää, E., McPhee, J. S., Narici, M. V., Gapeyeva, H., Pääsuke, M., Barnouin, Y., Hogrel, J.-Y., Butler-Browne, G., Bijlsma, A., et al. (2015). Plantarflexor Muscle-Tendon Properties are Associated With Mobility in Healthy Older Adults. J. Gerontol. A. Biol. Sci. Med. Sci. 70, 996–1002.

Suzuki, T., Bean, J. F. and Fielding, R. A. (2001). Muscle Power of the Ankle Flexors Predicts Functional Performance in Community-Dwelling Older Women. J.Am. Geriatr. Soc. 49, 1161–1167.

Thom, J. M., Morse, C. I.., Birch, K. M. and Narici, M. V. (2007). Influence of muscle architecture on the torque and power-velocity characteristics of young and elderly men. Eur. J. Appl. Physiol. 100, 613–619.

Wakahara, T., Kanehisa, H., Kawakami, Y., Fukunaga, T. and Yanai, T. (2013). Relationship between Muscle Architecture and Joint Performance during Concentric Contractions in Humans. J. Appl. Biomech. 29, 405–412.

Woodson, C., Bandy, W. D., Curis, D. and Baldwin, D. (1995). Relationship of Isokinetic Peak Torque With Work and Power for Ankle Plantar Flexion and Dorsiflexion. J. Orthop. Sports Phys. Ther. 22, 113–115.

Zellers, J. A., Carmont, M. R. and Silbernagel, K. G. (2018). Achilles Tendon Resting Angle Relates to Tendon Length and Function. Foot Ankle Int. 39, 343–348.

